# B cell receptor crosslinking can augment T cell help-mediated germinal center B cell selection

**DOI:** 10.1101/300939

**Authors:** Jackson S. Turner, Fang Ke, Irina L. Grigorova

**Author notes:** Please address correspondence to: Irina Grigorova, 1150 W. Medical Center Dr, 6748 MSII, Ann Arbor, MI 48109, (p), (734) (f).

## Abstract

Selection of germinal center (GC) B cells with B cell receptors (BCR) possessing high affinity to foreign antigen (Ag) and their differentiation into antibody-secreting long-lived plasma cells is critical for potent long-term humoral immunity. Ag-dependent engagement of GC B cell BCR triggers Ag internalization and loading of antigenic peptides on MHCII molecules for presentation to follicular helper T cells (Tfh) and acquisition of T cell help. However, whether it also provides signals that are critical or synergistic with T cell help for GC B cell selection and differentiation *in vivo* is not known. Here we show that T cell help is sufficient to induce GC B cell expansion and plasmablast (PB) formation in the absence of recurrent BCR engagement with Ag. Ag-mediated BCR crosslinking on the other hand is not sufficient to promote GC B cell selection, but can synergize with T cell help to enhance the GC B cell and PB responses when T cell help is limiting.

## Introduction

GCs are distinct sites within B cell follicles in which activated B cells undergo affinity maturation and can differentiate into memory B cells and long-lived plasma cells (Victora and Nussenzweig, 2012). The GC is characteristically polarized into the dark zone (DZ) and the light zone (LZ). The LZ contains Tfh cells and follicular dendritic cells (FDCs) on which antigen (Ag) is deposited for acquisition by GC B cells. GC B cells with higher affinity for Ag undergo selection in the LZ and then move into the DZ where they proliferate and undergo somatic hypermutation (Allen et al., 2007, Victora et al., 2010, Victora and Nussenzweig, 2012), Through ubiquitination and rapid degradation of MHCII-Ag complexes, they refresh their pool of MHCII molecules and reenter into the LZ for another round of Ag acquisition and presentation (Bannard et al., 2016, Victora et al., 2010, Mesin et al., 2016). Affinity-based selection of GC B cells is based on competition for Ag and T cell help, but the individual roles of the signals provided by BCR engagement and T-cells in promoting selection are not fully understood (Shlomchik and Weisel, 2012). Several studies have examined the effect of increasing T cell help to GC B cells independently of BCR cross-linking by taking advantage of the fact that GC B cells express high levels of DEC-205, a cell-surface lectin that delivers Ags it binds to MHCII loading compartments (Bonifaz et al., 2002, Victora et al., 2010). These studies found that upon administration of DEC-205 antibodies conjugated to T cell Ag (αDEC-Ag), high-affinity GC B cells deficient for DEC-205 were outcompeted by those that expressed the lectin. They also showed that administration of αDEC-Ag during GC responses increased the rate of GC B cell proliferation and hypermutation in a dose-dependent fashion, and promoted formation of PB (precursors of plasma cells) (Victora et al., 2010, Gitlin et al., 2014, Gitlin et al., 2015). These results suggest that competition for T cell help can drive GC B cell selection. However, recipient mice in these studies were immunized with B cell cognate Ag to initiate GC responses, raising the possibility that integration with signals from BCR engagement with Ag is necessary for αDEC-Ag-mediated enhancement of GC responses.

Indeed, recent evidence supports an important role for BCR engagement in promoting GC B cell selection and differentiation into PBs, despite the inhibition of the BCR signaling pathway in GC B cells (Khalil et al., 2012, Mueller et al., 2015, Nowosad et al., 2016). One study found that blocking GC B cells’ ability to acquire Ag inhibited initial PB differentiation more effectually than blockade of CD40 or depletion of Tfh cells (Krautler et al., 2017). Additionally, a recent study found that the signaling pathways downstream of both BCR and CD40, a critical ligand for mediating T cell help to GC B cells, are altered in GC B cells compared to naïve B cells such that stimulation through both is required for efficient induction of Myc, a critical driver of B cell proliferation (Luo et al., 2018). These findings suggest that although BCR signaling pathways are attenuated in GC B cells, they may nevertheless play a critical role in GC B cell selection and differentiation. However, the direct impact of Ag-dependent BCR engagement on GC B cells’ expansion or differentiation into antibody-secreting PBs could not be determined due to the difficulty of controlling GC B cells’ acquisition of Ag *in vivo*.

We recently described an experimental system that enables B cells to participate in GCs after a single transient acquisition of Ag (Turner et al., 2017c). Here we adapted it to recruit B cells into GCs and provide them with the means to acquire potential positive selection signals from BCR crosslinking or T cell help independently or in combination to examine the contributing roles of both signals to GC B cell survival, selection, and effector differentiation. We found that T cell help is sufficient to promote GC B cell expansion and PB differentiation in the absence of Ag-mediated BCR crosslinking. Conversely, Ag engagement of BCRs is insufficient to promote GC B cell selection in the absence of T cell help, but is able to synergistically enhance the GC B cell and PB responses when T cell help is limiting.

## Results and Discussion

To address the individual roles for antigen (Ag)-driven BCR cross-linking and T cell help in promoting GC B cell selection and effector differentiation, purified Hy10 Ig-Tg B cells (with BCRs specific to duck egg lysozyme, DEL) were first incubated *ex vivo* for 5 minutes with 50 μg/mL of DEL-OVA (DEL conjugated to ovalbumin, OVA), washed extensively, and transferred into recipient mice in which OTII Th cells were activated 3d before by immunization with OVA in complete Freund’s adjuvant (CFA) (**Fig. 1A**). As we have shown before, transient Ag acquisition *ex vivo* and cognate T cell help *in vivo* enable Ig-Tg B cells’ proliferation and participation in GCs, with recruitment into GCs starting by 4d post transfer (d.p.t) (**Fig. S1A-C**, (Turner et al., 2017c)). At this time point, due to the lack of cognate DEL Ag in the OVA-immunized recipient mice, Ig-Tg B cells should not receive any stimulation via Ag-dependent BCR cross-linking. In addition, prior to their differentiation into GC B cells, Ig-Tg cells undergo extensive proliferation (**Fig. S1C**), diluting the Ag peptides acquired during the *ex vivo* pulsing with DEL-OVA. Due to this dilution, the Ig-Tg B cells recruited into GCs are likely to present substantially lower amounts of OVA peptides than endogenous OVA-specific GC B cells, which can reacquire Ag *in vivo*. To summarize, by 4 d.p.t. Ig-Tg cells convert into GC B cells that are not subjected to Ag-dependent BCR crosslinking and should poorly compete for help from OVA-specific Tfh cells within GCs.

**Figure 1.**
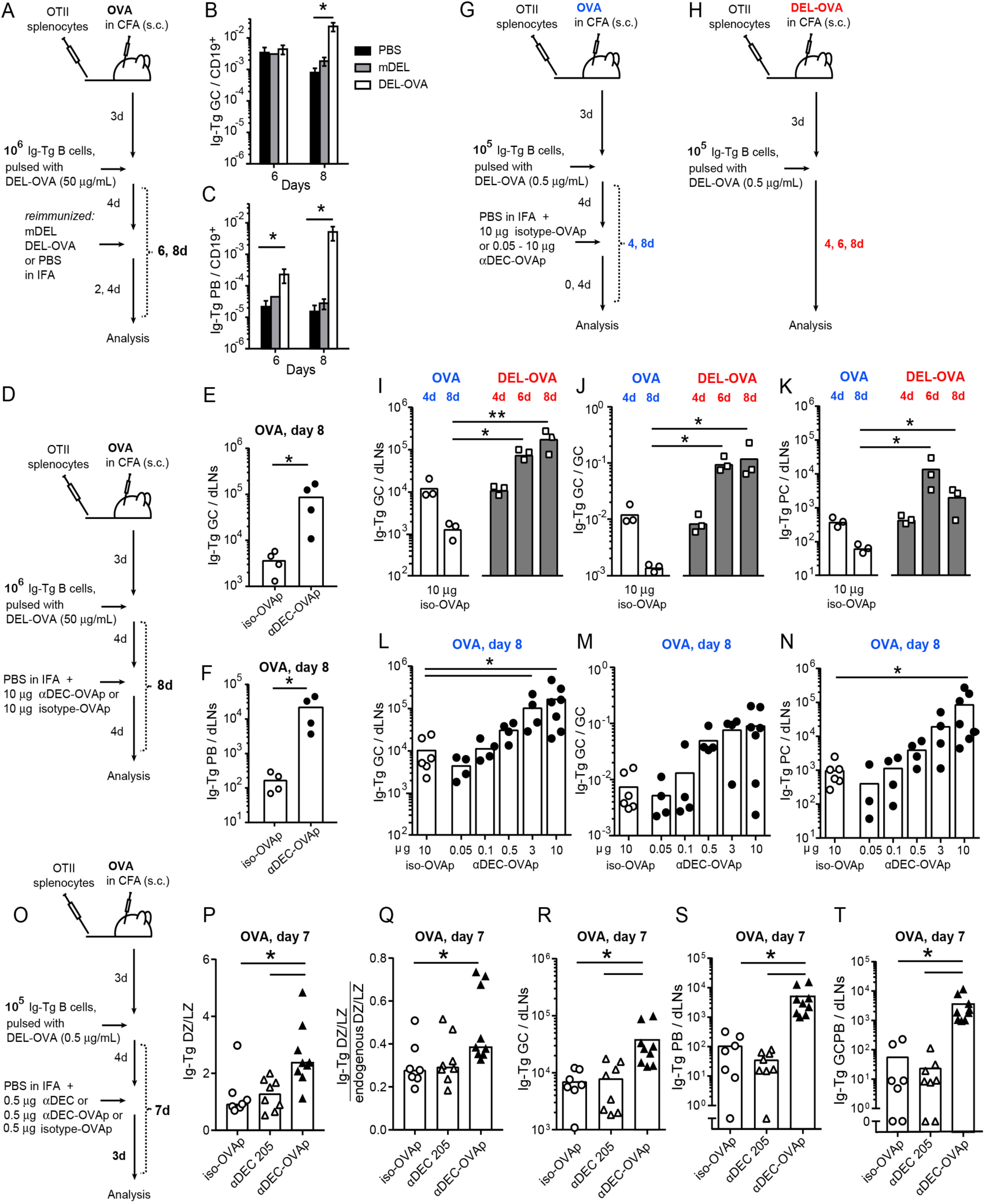
T cell help is sufficient for GC B cell selection and PB differentiation. **A**, Experimental outline for **B, C**. Purified Hy10 Ig-Tg B cells were pulsed *ex vivo* for 5 min with 50 μg/mL DEL-OVA, washed, and 10^6^ were transferred to recipient B6 mice preinjected with splenocytes containing 5×10^5^ OTII Th cells and subcutaneously (s.c.) preimmunized with OVA in CFA. Four days after Ig-Tg transfer, recipient mice were s.c. re-immunized with mDEL, DEL-OVA, or PBS in IFA. **B, C** Accumulation of Ig-Tg GC (**B**) and PBs (**C**) per CD19^+^ cells in the dLNs of re-immunized recipient mice at 2 and 4d post reimmunization (6 and 8d post Ig-Tg B cell transfer). See also **Fig. S1A-E**. **D,** Experimental outline for **E, F**. 10^6^ 50μg/ml DEL-OVA pulsed Ig-Tg B cells were recruited into GCs as in **A**, and 4 d.p.t. recipient mice were s.c. re-immunized with PBS in IFA and injected with 10μg of αDEC-OVAp or iso-OVAp. **E, F**, Ig-Tg GC (**E**) and PB (**F**) accumulation in dLNs from mice that received 10μg iso-OVAp (open symbols) or αDEC-OVAp (closed symbols) 4d earlier. See also **Fig. S1F**. **G**, Experimental outline for **I**-**N**. 10^5^ 0.5μg/ml DEL-OVA pulsed Ig-Tg B cells were recruited into GCs as in **A**, and 4 d.p.t. recipient mice were s.c. re-immunized with PBS in IFA and injected the indicated amount of αDEC-OVAp or iso-OVAp. **H**, Experimental outline for **I**-**K**. Recipient mice were injected with OTII splenocytes as in **A** and s.c. immunized with DEL-OVA in CFA. 10^5^ 0.5μg/ml DEL-OVA pulsed Ig-Tg B cells were transfered as in **G** and dLNs analyzed at indicated time points. **I, L**, Ig-Tg GC B cell accumulation in dLNs. **J, M,** Fraction of Ig-Tg GC B cells in GCs. **K, N**, Ig-Tg PB accumulation in dLNs. See also **Fig. S1G, H. O**, Experimental outline for **P-T**. Mice were treated as in **G** and at 4d post Ig-Tg cell transfer reimmunized with PBS in IFA and s.c. injected with 0.5μg αDEC-OVAp, iso-OVAp, or unconjugated αDEC-205 for analysis 3 days later (7 days after Ig-Tg cell transfer). **P**, Ig-Tg GC DZ to LZ ratio. **Q**, Ig-Tg DZ to LZ ratio normalized to the endogenous GC DZ to LZ cell ratio. **R**, Ig-Tg GC B cells accumulation in dLNs. **S**, **T**, Ig-Tg PB (**S**), and GCPB (**T**) accumulation in dLNs. See also **Fig. S1I-O. B, C,** Data from 3–5 independent experiments, 3–6 mice per condition, shown as mean ± SEM.* p<0.05, Kruskal-Wallis with Dunn’s post test between PBS, mDEL, or DEL-OVA. **E–T,** Data from 2–4 independent experiments. Each symbol represents one mouse. *, p<0.05, Mann-Whitney test (**E, F**) or Kruskal-Wallis with Dunn’s post test between isotype and each αDEC-OVAp dose (**I-N**), or between all conditions (**P-T**).

To address whether BCR cross-linking is sufficient to promote GC B cell expansion or the plasma cell response, at 4 d.p.t. of DEL-OVA-pulsed Ig-Tg B cells the recipient mice were re-immunized with 50 μg of multivalent DEL (mDEL) in incomplete Freund’s adjuvant (IFA) or with PBS in IFA for negative control (**Fig. 1A**). While mDEL could engage Ig-Tg GC B cells’ BCRs, they should not provide additional Ag peptides to present to OVA-specific Tfh cells. As positive controls, recipient mice received DEL-OVA in IFA to provide both additional BCR cross-linking of Ig-Tg GC B cells, as well as peptides to present to OVA-specific Tfh cells. Of note, in *ex vivo* stimulation assays, mDEL and DEL-OVA induce similar Ig-Tg BCR cross-linking and internalization (Turner et al., 2017). Draining inguinal LNs (dLNs) were collected 2 and 4d after re-immunization, and Ig-Tg GC B cells and PB were measured by flow cytometry (**Fig. 1A,** for gating see **Fig. S1A, D**). No increase in Ig-Tg GC or PB accumulation was detected after reimmunization of mice with mDEL compared to PBS control. However, a significant accumulation of Ig-Tg GC B cells and PBs was observed in DEL-OVA reimmunized recipients (**Fig. 1B, C, Fig. S1E**). These data suggest that elevated presentation of OVA peptides for acquisition of T cell help is necessary to promote Ig-Tg GC B cell selection and formation of PBs, while crosslinking of GC BCRs by itself is not sufficient.

While this finding is consistent with a well-established requirement of T cell help for GC response (Takahashi et al., 1998, Victora et al., 2010), it does not discriminate whether Ag-dependent BCR engagement is necessary for GC B cells in any other way than for deposition of Ag peptides on MHCII. A recent study suggested that combination of BCR signaling and T cell help may be required for GC B cell proliferation and selection (Luo et al., 2018). We therefore next asked whether T cell help was sufficient to promote GC B cell cycling and PB accumulation *in vivo*, or whether BCR cross-linking was also required. To address this question, we used αDEC-205 antibodies to target Ag to MHCII loading compartments in GC B cells, as previously described (Victora et al., 2010). Binding to the lectin DEC-205, which is upregulated on GC B cells, enables Ag to be loaded on MHCII without engaging the BCR (Victora et al., 2010). We conjugated the OVA peptide 323-339 (OVAp), which contains the OTII T cell epitope of OVA, to αDEC-205 or isotype control antibodies (αDEC-OVAp, iso-OVAp). DEL-OVA pulsed Ig-Tg B cells were recruited into GCs as above and at 4 d.p.t. recipient mice were reimmunized with PBS in IFA and s.c. injected with αDEC-OVAp or iso-OVAp to drain to inguinal LNs (**Fig. 1D**). Of note, both Ig-Tg and endogenous GC B cells express DEC-205 and would receive additional OVAp to present for T cell help upon administration of αDEC-OVAp. Significant increases in Ig-Tg GC B cell and PB accumulation were observed at 4d post administration of αDEC-OVAp, suggesting that elevated presentation of OVA peptides by GC B cells may be sufficient to induce their selection and differentiation in the absence of additional crosslinking of BCRs (**Fig. 1E, F, Fig. S1F**).

To verify that the observed results were not due to the transfer of small amounts of Ag by the DEL-OVA-pulsed Ig-Tg B cells *in vivo*, we significantly reduced the amount of Ag initially acquired by Ig-Tg B cells by pulsing them *ex vivo* with only 0.5μg/mL of DEL-OVA which is slightly above the threshold dose required for BCR-driven activation of Ig-Tg B cells (Turner et al., 2017c). To further minimize the potential for Ag transfer, the number of DEL-OVA pulsed Ig-Tg B cells transferred to recipient mice was reduced to 10^5^, of which the vast majority localizes to the spleens rather than peripheral LNs (Turner et al., 2017c). As before, DEL-OVA pulsed Ig-Tg B cells were extensively washed and transferred into OVA-immunized recipient mice in which activated OTII Th cells were present. 4 days later the recipients were reimmunized with PBS in IFA and injected with 0.05-10 μg of αDEC-OVAp or with 10μg iso-OVAp as negative controls (**Fig. 1G**). To compare the rescue of Ig-Tg GC and PB response by αDEC-OVAp administration to a more conventional immunization scenario, we also assessed the kinetics of Ig-Tg B cell participation in the GC and PB response in mice immunized with DEL-OVA, where transferred Ag-pulsed cells could reacquire cognate Ag *in vivo* (**Fig. 1H**). Draining LNs were collected at various times and Ig-Tg GC participation and PB accumulation were analyzed (**Fig. 1G, H**). As previously described, at 4 d.p.t. Ag-pulsed Ig-Tg B cells were similarly recruited into GC and PB responses in OVA or DEL-OVA immunized mice (**Fig. 1I-K**) (Turner et al., 2017b, Turner et al., 2017c). However, 4 days later Ig-Tg B cells started to drop out of GCs in the OVA-immunized mice. In contrast, in DEL-OVA immunized mice they expanded to around 10% of total GC B cells (**Fig. I, J**). Ig-Tg PB production was maximized in DEL-OVA immunized mice at 6 d.p.t. (**Fig. 1K**). In the OVA-immunized mice (**Fig. 1G**) administration of αDEC-OVAp led to dose-dependent rescue of Ig-Tg GC and PB responses (**Fig. 1L-N**). Ig-Tg B cell fraction in GCs was maximized at around 10% at 3 μg of αDEC-OVAp (**Fig. 1L, M**). Of note, Tfh cell response was largely consistent across various amounts of αDEC-OVAp administered, except of the highest 10 μg dose of αDEC-OVAp when a trend towards a larger Tfh cell population was observed (**Fig. S1G, H**). To control for potential effects of αDEC-205 ligation independent of Ag loading, we analyzed the Ig-Tg GC and PB response at 3 days after the administration of the intermediate 0.5 μg dose of αDEC-OVAp compared to the iso-OVAp or unconjugated αDEC205 negative controls (**Fig. 1O**). As expected, only when conjugated to OVAp, αDEC205 induced augmented cycling of Ig-Tg GC B cells as based on the ratio of the dark zone (DZ) to light zone (LZ) GC B cells (**Fig. S1I, Fig. 1P, Q**) and an overall increase in the Ig-Tg GC response (**Fig. 1R**), while the endogenous GC response was not affected (**Fig. S1L**). Of note, at 2 d post αDEC-OVAp administration Ig-Tg B cell cycling was not yet elevated (**Fig. S1P, Q**), presumably due to some time required for Ig-Tg cells to integrate T cell help when OVAp peptide is loaded through DEC205 binding onto both Ig-Tg and endogenous GC B cells. To analyze both the Ig-Tg and endogenous PB responses we performed intracellular staining with anti pan-Ig, which yields similar numbers for Ig-Tg PBs as intracellular HEL staining and enables quantification of endogenous PBs (**Fig. S1D, J, K).** In addition, by gating on the Fas^+^ CD38^lo^ Ig^high^ cells, we identified the PB that recently originated from GCs (GCPB, **Fig. S1O**, (Krautler et al., 2017)). We found that αDEC-OVAp administration leads to an increase in the total Ig-Tg PB, as well as GCPB response, while the endogenous PB response is not significantly elevated (**Fig. 1S, T, Fig. S1M, N**). Given that DEC205 is expressed on both endogenous and Ig-Tg GC B cells, the selective increase in Ig-Tg GC and GCPB responses following αDEC-OVAp administration indicates that this treatment narrows the difference in the amount of OVAp presented by Ig-Tg and endogenous GC B cells, enabling them to more successfully compete for T cell help over time (**Fig. 1G-N**). Altogether these results indicate that GC and PB responses are enhanced when GC B cells receive additional sources of peptide to present for T cell help, and suggest that Ag-dependent BCR engagement is not absolutely required to promote GC B cell expansion or PB formation.

We then addressed whether Ag-mediated BCR cross-linking may be able to enhance GC B cell selection or PB differentiation in combination with saturating or sub-saturating amounts of T cell help. To test that, Ig-Tg B cells were recruited into GCs as above, and the recipient mice were injected with 0.5μg or 10μg αDEC-OVAp and reimmunized with PBS or 50 μg mDEL in IFA. As negative controls, mice received isotype-OVAp (**Fig. 2A**). Draining LNs were analyzed 3d later. The Ig-Tg GC and PB responses were not enhanced following re-immunization with mDEL compared to PBS in recipients that received either isotype-OVAp or 10μg αDEC-OVAp, suggesting that BCR crosslinking does not promote GC B cell selection in the absence of T cell help or in the presence of saturating T cell help (**Fig. 2B-E, J-M**).Interestingly, in the absence of additional peptide presentation for Tfh cells, administration of 50 μg of mDEL led to modest reduction in Ig-Tg GC B cells (**Fig. 2C,** isotype-OVAp), which was reversed when GC B cells were presenting some cognate peptides for T cell help (**Fig. 2G, K**, αDEC-OVAp). The observed effect is consistent with previous studies where acute administration of a large dose of cognate to BCR Ag (or membrane-associated presentation of cognate self-Ag in proximity to GCs) reduced the numbers of GC B cells (Pulendran et al., 1995, Shokat and Goodnow, 1995, Silva et al., 2017, Chan et al., 2012). Based on the previous studies and our work (Turner et al., 2017a), we speculate that receiving BCR signaling in the absence of sufficient T cell help may purge GC B cells from prolonged “hanging on” participation in GCs.

**Figure 2.**
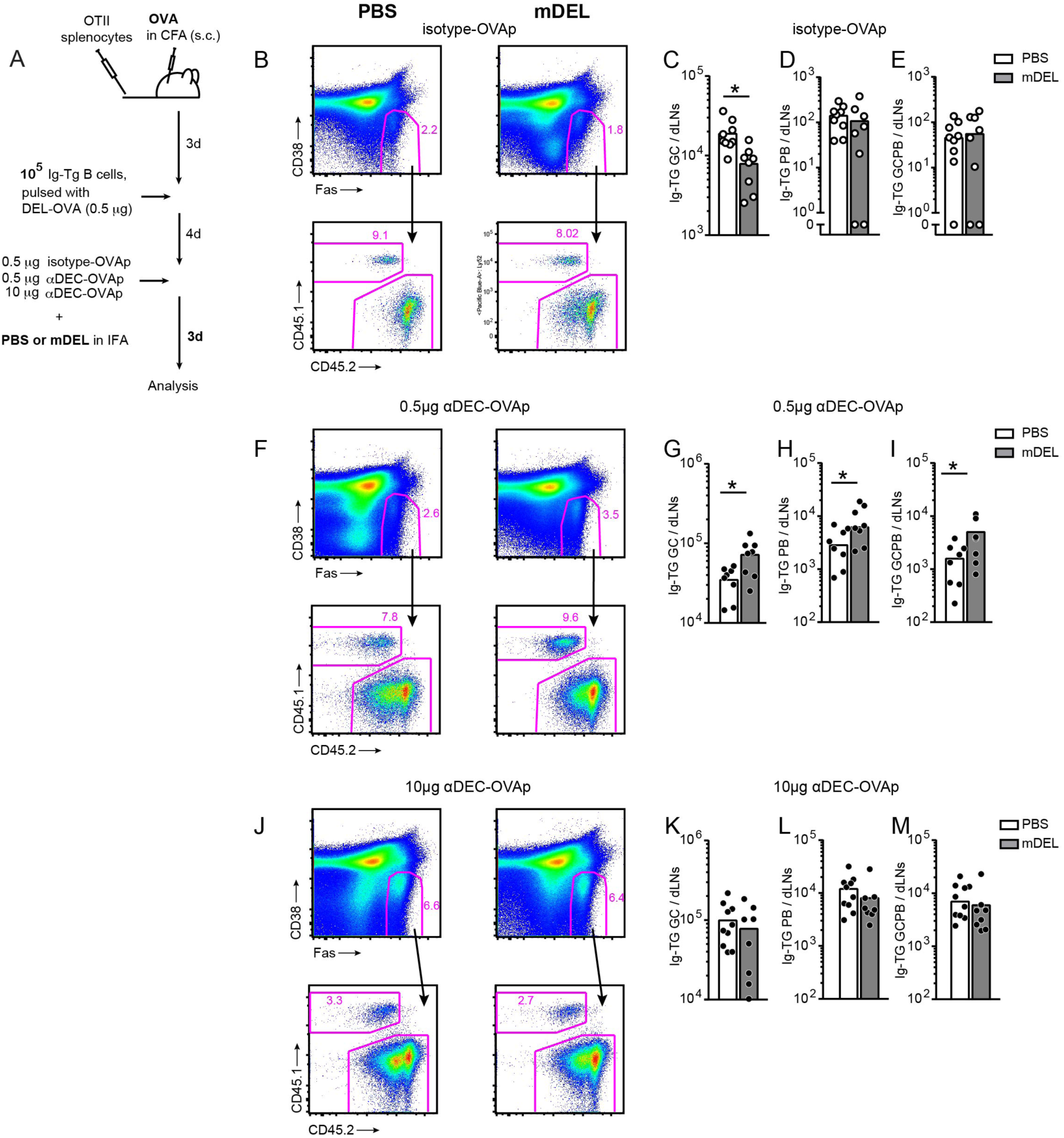
BCR Cross-linking promotes GC and PB response when T cell help is sub-saturating. **A**, Experimental outline. 0.5 μg/mL DEL-OVA pulsed Ig-Tg B cells were recruited into GC responses as in **Fig. 1G**. 4 d.p.t., recipient mice were s.c. re-immunized with PBS or 50 μg mDEL in IFA and injected with 0.5μg iso-OVAp or with 0.5μg or 10μg αDEC-OVAp. **B-K**, Representative examples of GC B cell gating analysis of Ig^+^ CD3^−^ lymphocytes in dLNs at 3 days post reimmunization (7 days post Ig-Tg cell transfer) (in **B, F, J**) and accumulation of Ig-Tg GC B cells (**C, G, K**), PB (**D, H, L**) and GCPB (**E, I, M**) in dLNs. Data from 2–4 independent experiments. Each symbol represents one mouse. *, p<0.05, Mann-Whitney test. See also **Fig. S2.**

In contrast, after administration of an intermediate dose of αDEC-OVAp (0.5μg) that by itself led to only a modest increase in Ig-Tg responses (**Fig. 1L, N**), re-immunization with mDEL in IFA led to accumulation of Ig-Tg GC and PB compared to PBS in IFA control (**Fig. 2F-I)**. Reimmunization with 10 fold lower dose of mDEL (5 μg) also led to robust increases in the Ig-Tg GC, GCPB and PB responses (**Fig. S2A-F**). Finally, similar trends were obtained when mice were reimmunized with highly multivalent polystyrene microspheres coated with DEL (sphDEL) in IFA (**Fig. S2G-J**). These results suggest that Ag-dependent BCR engagement in GC B cells, while neither sufficient nor necessary for GC B cell selection, can enhance GC and PB responses in combination with T cell help.

Our findings are consistent with an *ex vivo* observation that simultaneous engagement of GC BCRs and CD40 enhances upregulation of Myc in GC B cells which is required for cell cycling (Luo et al., 2018). However, they suggest that T cell help may be sufficient on its own. Although CD40 is a critical component of the ‘help’ provided by Tfh cells, other T-cell derived factors may enable T cell help to promote expansion of GC B cells and formation of PBs independently of re-engagement of their BCRs with Ag. The importance of T cell help-mediated factors other than CD40 in promoting GC PB response was also suggested by another study in which depletion of CD4 T cells during the GC response inhibited PB response more profoundly than blockade of CD40L (Krautler et al., 2017).

Interestingly, the study by Krautler *et al.* also demonstrated that differentiation of PB from GC B cells was more effectively impeded by blockade of B cells’ acquisition of Ag than by depletion of T cells, suggesting that Ag-dependent BCR cross-linking may play a critical role in initiating GC B cells’ differentiation into PBs. However, whether BCR crosslinking alone may be sufficient to initiate formation of the early PB in the absence of T cell help has not been explored. We sought to address this question directly using our experimental system to induce BCR crosslinking and T cell help independently.

Early PBs differentiating from GC B cells were previously identified as having a GC phenotype (CD38^lo^) and intermediate expression of the transcription factor Blimp1 and surface Ig, whereas later PBs had higher Blimp1 expression and lower surface Ig. Additional characterization of these populations indicated that early PBs had increased surface expression of B220 and CD45 compared to more mature PBs (Krautler et al., 2017). Using Blimp1 reporter Ig-Tg B cells, we identified that B220 downregulation could be used as a surrogate marker of PB and GCPB maturation, as B220^+^ GCPBs expressed lower amounts of Blimp1 and syndecan, and higher amounts of CD86, surface IgG_1_, and CD45.1 than their B220^lo^ counterparts (**Fig. 3A**, **B, Fig. S3A**). To determine whether BCR cross-linking is sufficient to promote early differentiation of PBs in GC B cells, Ig-Tg B cells were recruited into the GC response as above, and recipient mice were injected with 0.5μg isotype-OVAp or αDEC-OVAp. The recipient mice were then reimmunized with PBS, mDEL, or sphDEL in IFA, and the GC PB response was measured 1 and 2d later (**Fig. 3C, D**). We found that BCR crosslinking alone was insufficient to increase accumulation of early GCPBs, independently of whether moderately multivalent Ag mDEL or highly multivalent sphDEL were used for reimmunization (**Fig. 3E-G**). Of note, when analysis of early PB differentiation was confined to the IgG_1_^+^-cells we detected a potential increase in the early Ig-Tg PBs at 2d following reimmunization with highly multivalent sphDEL (**Fig. S3B**). If confirmed, these results could indicate an isotype-dependent sensitivity of GC B cells to highly multivalent Ag. Alternatively, the increase in these cells could represent increased class switching to IgG_1_ in response to highly multivalent Ag. Finally, at 1-2 days after reimmunization we detected no increase in the total or DZ Ig-Tg GC B cells in response to BCR crosslinking alone, independently of their class-switching to IgG_1_ (**Fig. 3H, Fig. S3C, D, data not shown**).

**Figure 3.**
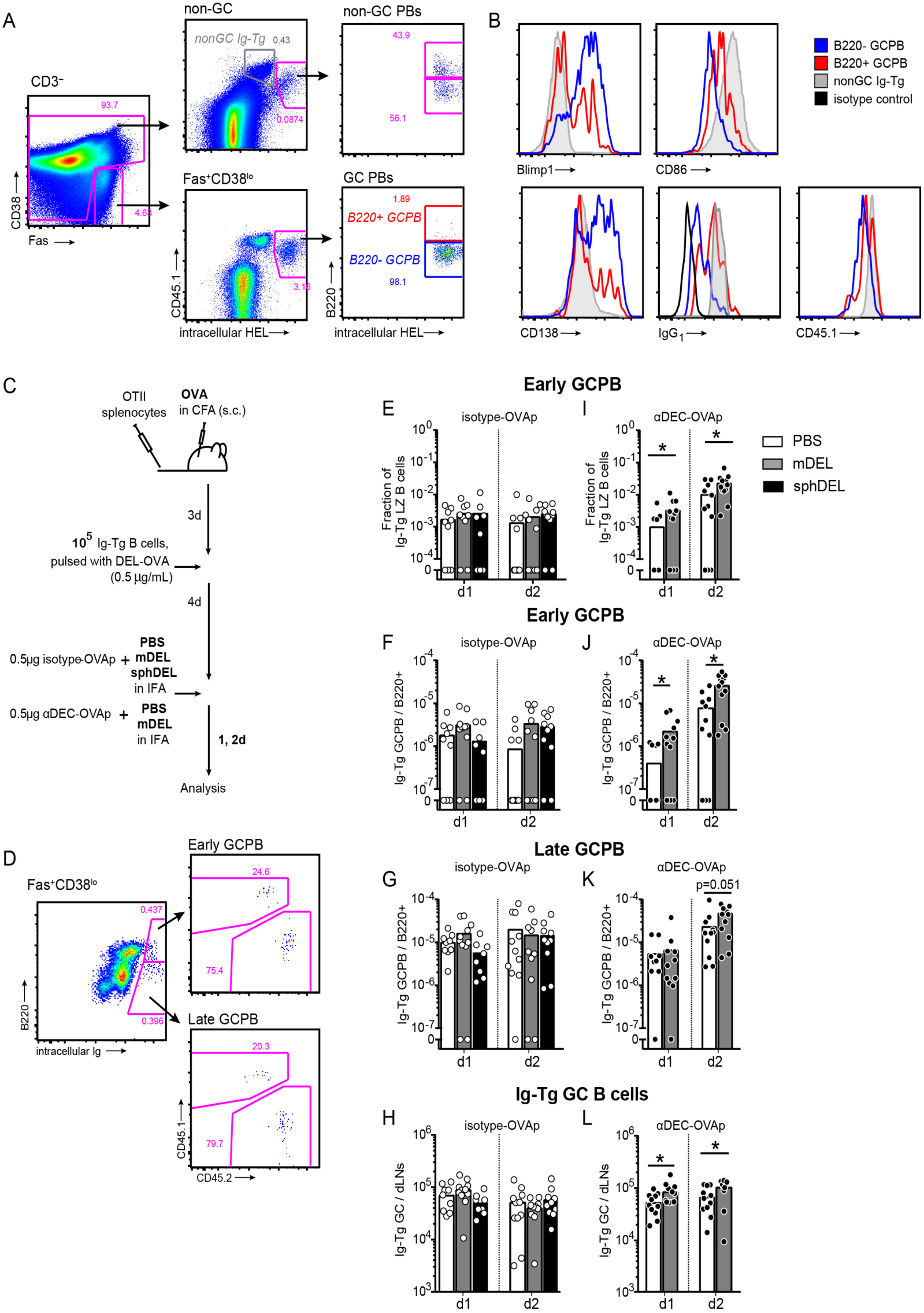
BCR cross-linking enhances early PB differentiation when T cell help is sub-saturating. **A**, Gating strategy for Ig-Tg early and late PBs and GCPBs. 10^5^ Blimp1^yfp^ or regular CD45.1^pos^ Ig-Tg B cells were transferred to B6 recipient mice, which were immunized s.c. with 50μg DEL-OVA. dLNs were analyzed 10 d.p.i. **B**, representative histographs of Blimp1, CD86, CD138, surface IgG_1_, and CD45.1 in B220^+^(red) and B220^lo^ (blue) GCPBs and non-GC Ig-Tg B cells (grey). For IgG_1_ staining non-GC B cells were gated on IgG_1_^pos^ cells. Data representative of 2-3 independent experiments with 13 mice. **C**, Experimental outline for **D-L**. 0.5 μg/mL DEL-OVA pulsed Ig-Tg B cells were recruited into GC responses as in **Fig. 1G**. Four d.p.t., recipient mice were s.c. re-immunized with PBS, 50 μg mDEL or sphDEL in IFA and injected with 0.5μg iso-OVAp or αDEC-OVAp. **D**, Representative gating example for Ig-Tg and endogenous B220^+^ early GCPBs and B220^lo^ late GCPBs. **E-L**, accumulation of B220^+^ early (**E-J**) and B220^lo^ late (**G, K)** GCPBs and GC B cells **(H, L)** in dLNs 1d and 2d post reimmunization. Data from 4 independent experiments. Each symbol represents one mouse. *, p<0.05, Kruskal-Wallis with Dunn’s post test between PBS and mDEL or sphDEL (left column), or Mann-Whitney test. See also **Fig. S3.**

In contrast, in the presence of subsaturating amounts of αDEC205-OVAp, BCR crosslinking promoted Ig-Tg early GCPB responses, as well as Ig-Tg GC B cell expansion and accumulation in the DZ (**Fig. 3I-L, Fig. S3E-G**). At the same time no accumulation of non-GCPB has been observed (**Fig. 3A**, **Fig. S3H-K**). Therefore, in the presence of subsaturating T cell help, Ag-dependent BCR crosslinking can promote increased differentiation of GC B cells into PBs and induce an increase in GC B cell expansion. Overall, our findings are consistent with the previously discovered role of Ag-dependent BCR engagement in driving GC differentiation into PB. However, while in the previous study T cell help has been shown to be critical for maturation and survival of the early GCPB (Krautler et al., 2017), our findings are more consistent with another study (Ise et al., 2018) and suggest that an ongoing T cell help is required, even for the initial differentiation of GC B cells into PBs.

To summarize, in this study we found that BCR crosslinking is not sufficient to promote GC B cell expansion, selection or differentiation into PB. In contrast, acquisition of T cell help is sufficient to induce GC B cell expansion and PB formation even in the absence of BCR engagement with Ag (**Fig. 4**). These findings are consistent with a recent study which showed that in the presence of abundant T cell help, non-Ag specific B cells could participate in GCs and persist long enough to acquire specificity to Ag (Silver et al., 2018).

**Figure 4.**
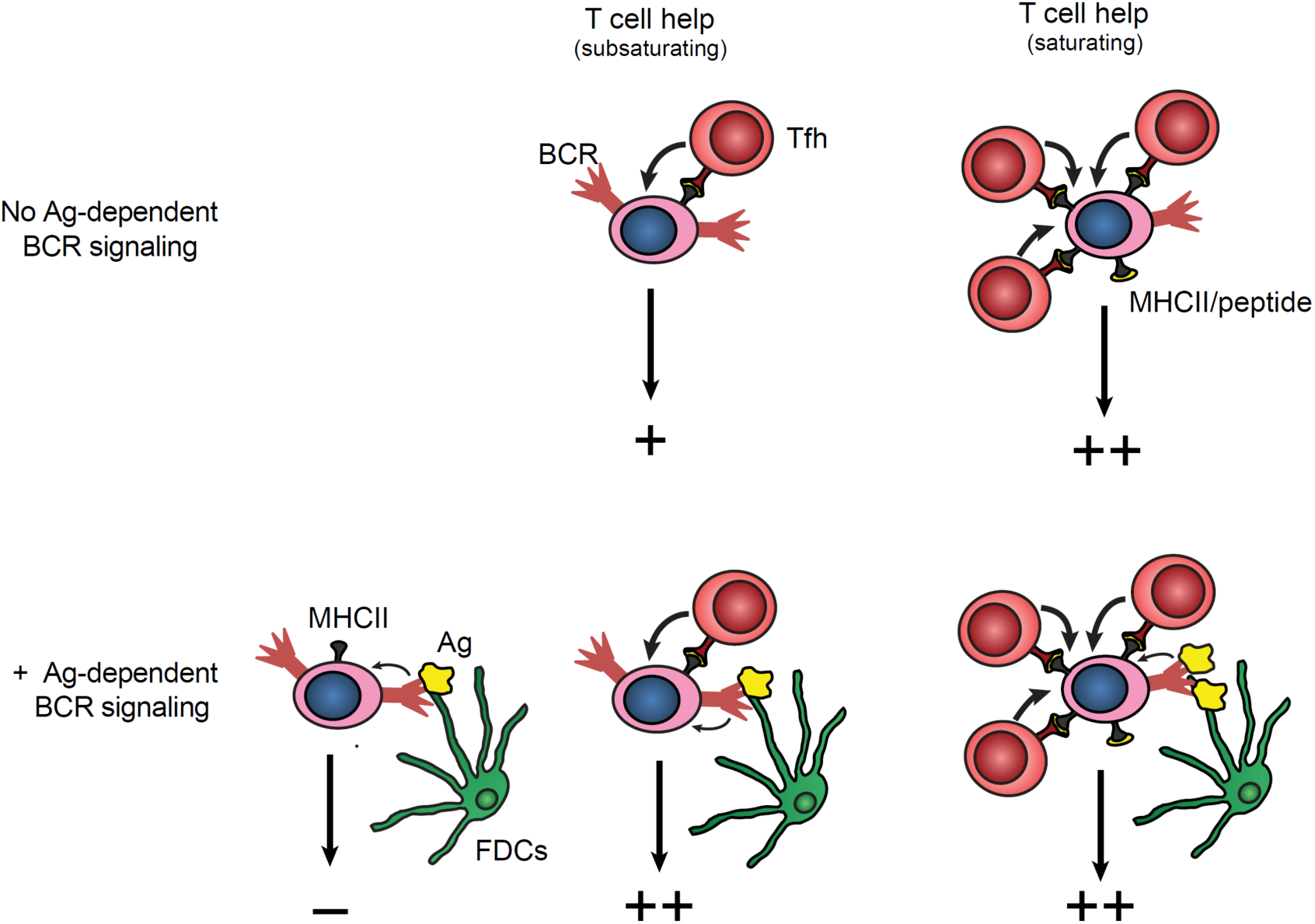
Model of GC B cell selection.

Although loading GC B cells with T cell Ag peptides was sufficient to promote their selection and PB differentiation in a dose-dependent fashion (**Fig. 1G, H,** (Gitlin et al., 2014)), we found that Ag-dependent BCR engagement potentiated GC B cell expansion and PB differentiation when the amount of T cell Ag peptide loaded was subsaturating (**Fig. 4**). The dynamics of GC B cells’ acquisition of Ag and T cell help *in vivo* are not completely understood. It is possible that when Ag is abundant, LZ GC B cells may reacquire Ag multiple times from FDCs and make serial productive contacts with Tfh cells that efficiently induce Myc and promote selection and differentiation. However, when Ag becomes more limiting, BCR signaling induced by Ag engagement is likely to be temporally separated from T cell help due to the time required for Ag digestion, MHCII loading and efficient competition with other GC B cells for T cell help. While BCR signaling was shown to induce transient nuclear exclusion of Foxo1 (a suppressor of Myc) in GC B cells *ex vivo*, it starts to decline 20 minutes after BCR engagement (Luo et al., 2018). Therefore, GC B cells are likely to have a very short window of time following Ag acquisition in which they need to acquire T cell help to synergistically induce Myc. Sufficiency of T cell help for GC B cell selection relieves this time constraint and ensures that GC B cell selection can continue when Ag reacquisition from FDCs becomes more limiting. While CD40 induces only minor upregulation of Myc *ex vivo*, additional factors produced by Tfh cells, such as BAFF, ICOS, and IL-21 may be sufficient to promote GC B cell selection and effector differentiation *in vivo* (Han et al., 1995, Ding et al., 2013, Goenka et al., 2014). Other factors promoting GC B cells’ survival or differentiation can also be provided by FDCs (e.g. BAFF, IL-6, complement fragments, and adhesion molecules) (El Shikh et al., 2010, Victoratos et al., 2006) or toll-like receptor ligands and could synergize with T cell help in the absence of Ag-dependent BCR engagement (Wang et al., 2011, Garin et al., 2010, Rookhuizen and DeFranco, 2014).

Overall, our current study suggests a dual role of BCR signaling and T cell help for GC B cell response *in vivo* with T cell help playing the dominant role and Ag-dependent BCR crosslinking enhancing GC B cell selection and differentiation into plasma cells (**Fig. 4**).

## Materials and Methods

### Mice

B6 (C57BL/6) and B6-CD45.1 (Ptprc^a^ Pepc^b^/BoyJ) mice were purchased from Charles River or the Jackson Laboratory. Blimp1^yfp^ (Prdm1-EYFP) mice were purchased from the Jackson Laboratory. BCR transgenic (Ig-Tg) Hy10 mice and TCR transgenic OTII mice were generously provided by Jason Cyster (Allen et al., 2007, Barnden et al., 1998). Hy10 mice were crossed with B6-CD45.1 and Blimp1^yfp^ mice. All mice were maintained in specific pathogen free environments and protocols were approved by the Institutional Animal Care and Use Committee of the University of Michigan.

### Antigen preparation and antibody conjugation

Duck eggs were locally purchased and lysozyme was purified as previously described (Allen et al., 2007). Ovalbumin (OVA) was purchased from Sigma. Duck egg lysozyme (DEL) was conjugated to OVA via glutaraldehyde cross-linking as previously described (Allen et al., 2007). For production of multimeric DEL (mDEL), purified DEL was conjugated to biotin at a 1:2 molar ratio using biotin-X NHS-ester (Pierce) according to the manufacturer’s directions and incubated with purified streptavidin (Thermo Scientific) at a 10:1 molar ratio for 30 minutes on ice, followed by removal of unbound DEL-bio by passage through a 30 kDa molecular weight cut-off desalting column (Bio-Rad).

For generation of DEL-coated microspheres (sphDEL), 0.11 μm streptavidin coated polystyrene microspheres (Bangs Laboratories) were diluted into PBS and combined with a saturating amount of DEL-bio as described previously (Eckl-Dorna and Batista, 2009)

For generation of αDEC-OVAp and iso-OVAp, purified DEC-205 (NLDC-145) and rat IgG2a isotype control antibodies were purchased from Biolegend and partially reduced with 50 mM 2-mercaptoethylamine (2-MEA) in PBS, 10 mM EDTA for 90 minutes at 37° C. 2-MEA was removed by passage through 30 kDa molecular weight cut-off desalting columns (Bio-Rad), and half-IgGs were incubated with a 9-fold molar excess of maleimide-substituted OVA peptide 323-339 (Genscript) for 2h at 4° C. Unbound peptide was removed by passage through 30 kDa molecular weight cut-off desalting columns (Bio-Rad) and conjugation was verified by SDS-PAGE.

### Adoptive transfer and immunization

Spleens were harvested from male donor OTII mice and pressed through 70μm nylon cell strainers (Falcon) in DMEM (Cellgro) supplemented with 2% FBS (Atlanta Biologicals), 10 mM HEPES, 50 IU/mL of penicillin, and 50 μg/mL of streptomycin (HyClone). Splenocytes were centrifuged for 7 minutes at 380 rcf, 4° C and resuspended in 0.14 M NH_4_Cl in 0.017 M Tris buffer, pH 7.2 for erythrocyte lysis, washed twice with DMEM supplemented as above, and counted using a Cellometer Auto X4 (Nexcelom). The fraction of CD19^−^ CD8^−^ CD4^+^ Vβ5^+^ (OTII) splenocytes was determined by flow cytometry, and the indicated number of OTII cells were transferred i.v. to male recipient mice. Ig-Tg B cells were enriched from male or female donor mice by negative selection as previously described (Allen et al., 2007). For transient exposure to Ag, purified Ig-Tg B cells were incubated with the indicated concentration of DEL-OVA *ex vivo* for 5 minutes at 37° C, washed four times with DMEM supplemented as above, and transferred i.v. to recipient mice. Where indicated, recipient mice were immunized s.c. in the flanks and base of tail with 50 or 5 μg of the indicated Ag emulsified in complete or incomplete Freund’s adjuvant (Sigma), prepared according to the manufacturer’s directions. Where indicated, recipient mice were injected s.c. in the base of tail with unconjugated αDEC-205, αDEC-OVAp or iso-OVAp in PBS.

### Flow cytometery

The following antibodies specific to B220 (RA3-6B2), CD19 (1D3), CD95 (Jo2), IgG1 (A85-1), IgG1a (10.9), rat IgG1 isotype control (R3-34) from BD-Pharmingen; CD4 (RM-45), CD45.1 (A20), CD45.2 (104), CD86 (GL-1), IgD (11-26c.2a), CD3 (17A2), CD197 (4B12), CD279 (RMP1-30), TCR Vβ5 (MR9-4) from Biolegend; CD-8 (53-6.7), CXCR5 (2G8), CD138 (281.2) from BD Bioscience, CD38 (90), GL-7 (GL-7), CXCR4 (2B11) from eBioscience have been used for flow cytometry analysis. Single-cell suspensions from draining lymph nodes were incubated with biotinylated antibodies for 20 minutes on ice, washed twice with 200 μl FACS buffer (2% FBS, 1mM EDTA, 0.1% NaN_3_ in PBS), incubated with fluorophore-conjugated antibodies and streptavidin (SA-Qdot 605, SA-Alexa647 from Life technologies; SA-Dylight 488 from Biolegends) for 20 minutes on ice, washed twice more with 200 μl FACS buffer, and resuspended in FACS buffer for acquisition. For intracellular staining, surface-stained cells were fixed and permeabilized for 20 minutes on ice with BD Cytofix/Cytoperm buffer, washed twice with 200 μl BD Perm/Wash buffer, incubated with Alexa 647-conjugated HEL or polyclonal goat anti-Ig(H+L) from Southern Biotech for 20 minutes on ice, followed by two more washes with 200 μl Perm/Wash buffer, and resuspended in FACS buffer for acquisition. Cells were acquired on a FACSCanto, and data was analyzed using FlowJo (TreeStar).

### Statistics

Statistical tests were performed as indicated using Prism 7 (GraphPad). Differences between groups not annotated by an asterisk did not reach statistical significance. No blinding or randomization was performed for animal experiments, and no animals or samples were excluded from analysis.

## Supplementary Figures

**Figure S1.**
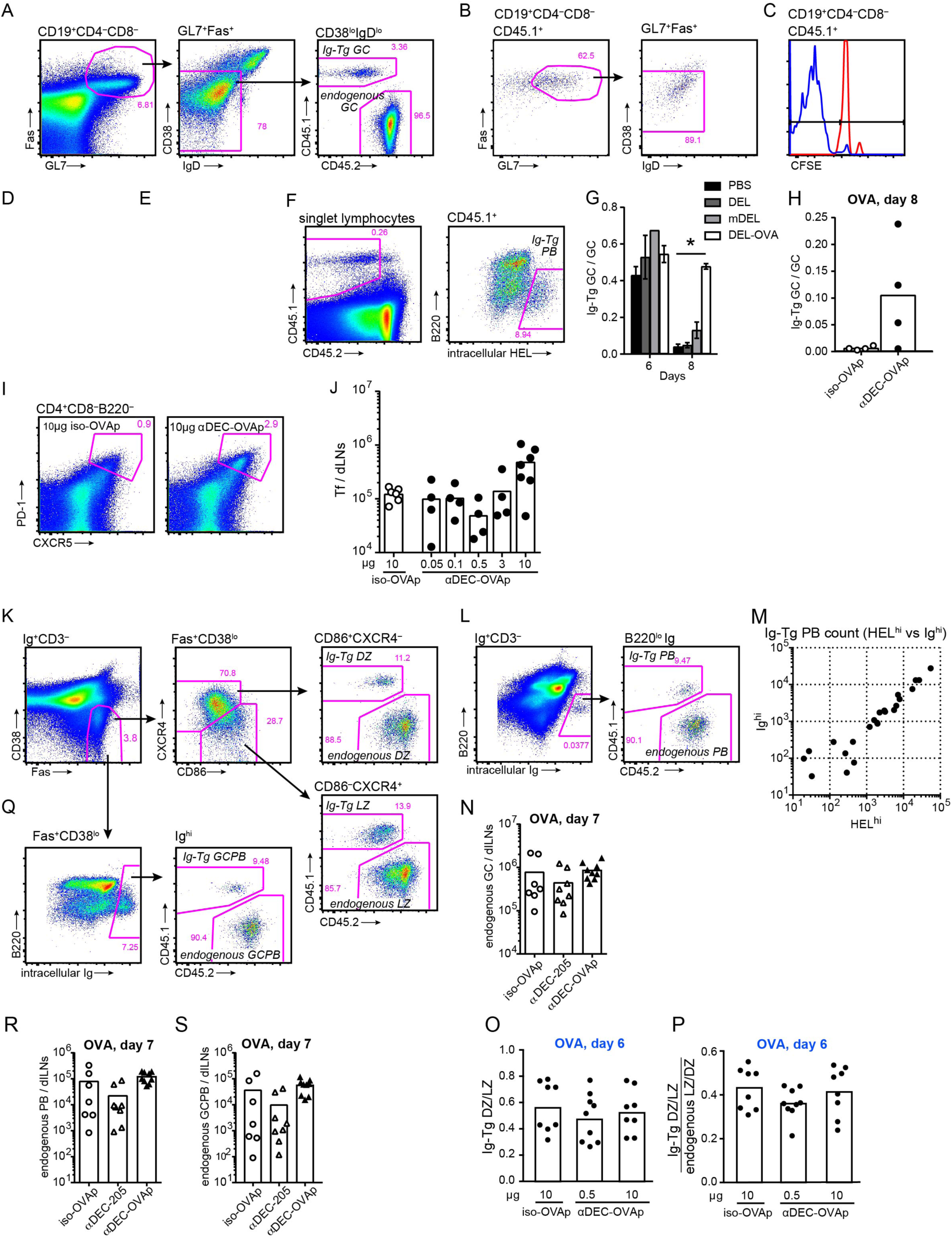
T cell help is sufficient for GC B cell selection and PB differentiation. **A**, **D**, Gating strategy for Ig-Tg and endogenous GC B cells (**A**) and Ig-Tg PBs (**D**). **B**, **C**, Representative plots of Ig-Tg GC participation (**B**) and proliferation (**C**) 4 d.p.t. according to the experimental design in **Fig. 1A**. Red line (ctrl) in **C** is from DEL-OVA pulsed Ig-Tg B cells transferred into unimmunized recipient mice. Representative of 3–5 independent experiments, 3–6 mice per condition. **E, F,** Ig-Tg GC B cells fraction of total GC cells analysis of the experiments shown in **Fig. 1A, B** (in **E**) and **Fig. 1D, E** (in **F**). **G, H**, Representative flow plots (**G**) and enumeration (**H**) of follicular CD4 T cells in dLNs of mice reimmunized with PBS in IFA and injected with 10μg iso-OVAp and αDEC-OVAp 4d earlier, according to the experimental design in **Fig. 1G**. **I, J, O,** gating strategies for Ig-Tg and endogenous DZ and LZ GC B cells (**I**), PB (**L**) and GCPB (**O**). **K**, Numbers of Ig-Tg PBs as determined by intracellular HEL and Ig staining recovered from dLNs of mice treated according to the experimental design in **Fig. 1G**. Data from 2 independent experiments. Each symbol represents one mouse. **L-N**, Endogenous GC (**L**), PB (**M**), and GCPB (**N**) accumulation in dLNs of recipient mice reimmunized with PBS and injected with 0.5μg iso-OVAp (open circles), unconjugated αDEC-205 (open triangles), or αDEC-OVAp (closed triangles) 3d earlier according to the experimental design in **Fig. 1O**. Data from 4 independent experiments. Each symbol represents one mouse. Kruskal-Wallis with Dunn’s post test between all conditions. **P, Q,** Analysis of Ig-Tg GC DZ to LZ cell ratio (in **P**) and Ig-Tg DZ/LZ ratio normalized to endogenous DZ/LZ GC B cell ratio (in **Q**) in experiment performed according to the experimental scheme in **Fig. 1G** and analyzed at 2d post administration of iso-OVAp and αDEC-OVAp (or 6 days post Ig-Tg B cell transfer).

**Figure S2.**
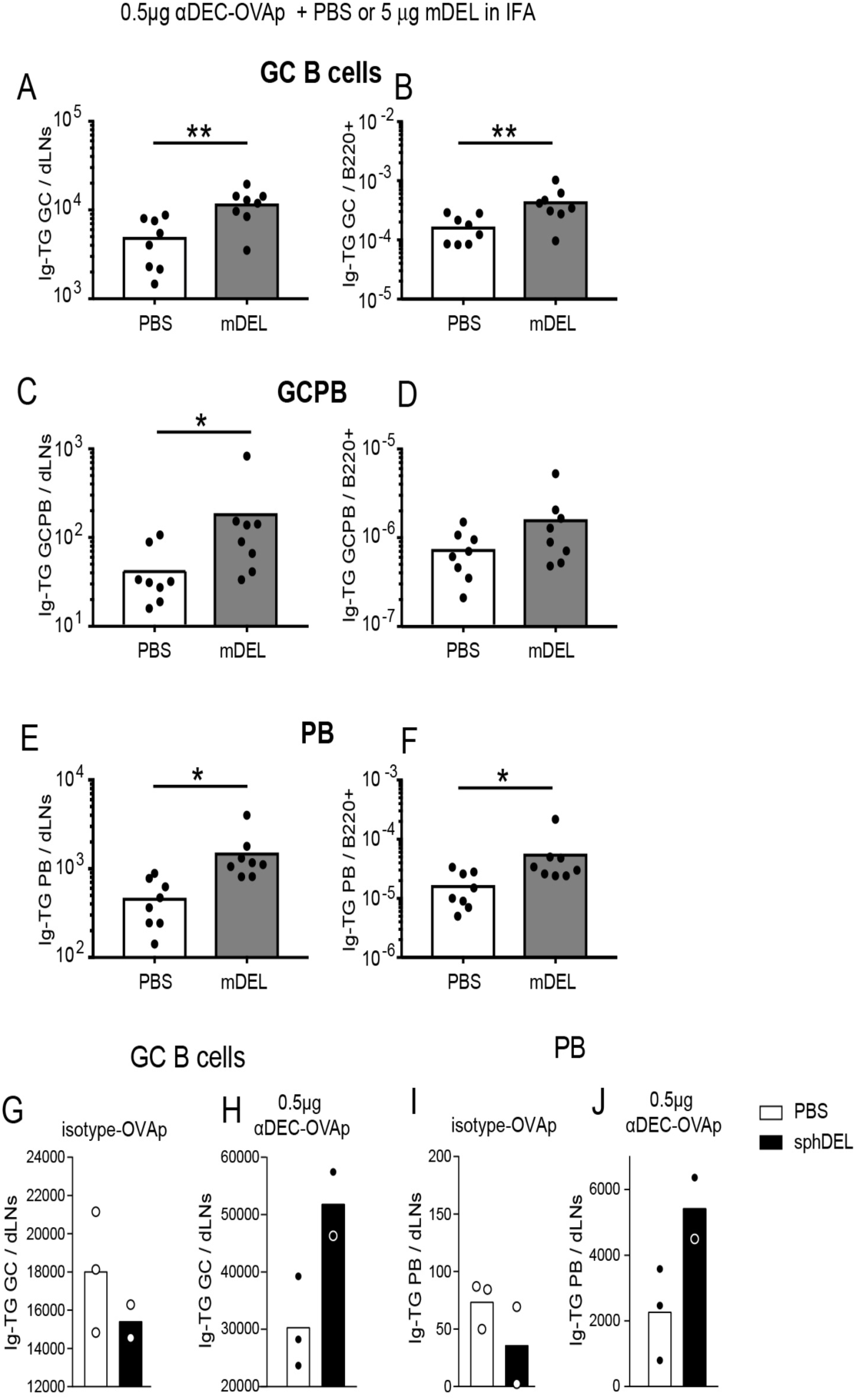
BCR Cross-linking promotes GC and PB response when T cell help is sub-saturating. **A-F**, Experimental design is similar to that in **Fig. 2A**, with recipient mice injected with 0.5μg αDEC-OVAp and s.c. reimmunized with PBS or 5 μg mDEL in IFA. Analysis of Ig-Tg GCs (**A, B**), GCPB (**C, D**) and PB (**E, F**) accumulation in dLNs (**A, C, E**) and per B220^+^ cells (**B, D, F**). Data from 2 independent experiments. Each symbol represents one mouse. *, p<0.05, **, p<0.01, Mann-Whitney test. **G-J**, Experimental design as in **Fig. 2A**, with recipient mice s.c. reimmunized with PBS or sphDEL in IFA and injected with 0.5μg iso-OVAp or αDEC-OVAp. Ig-Tg GC (**G, H**) and PB (**I, J**) accumulation in dLNs from mice reimmunized with PBS (white bars) or sphDEL (black bars). Data from 2–3 independent experiments, with 4–8 mice per condition. Each symbol represents one experiment.

**Figure S3.**
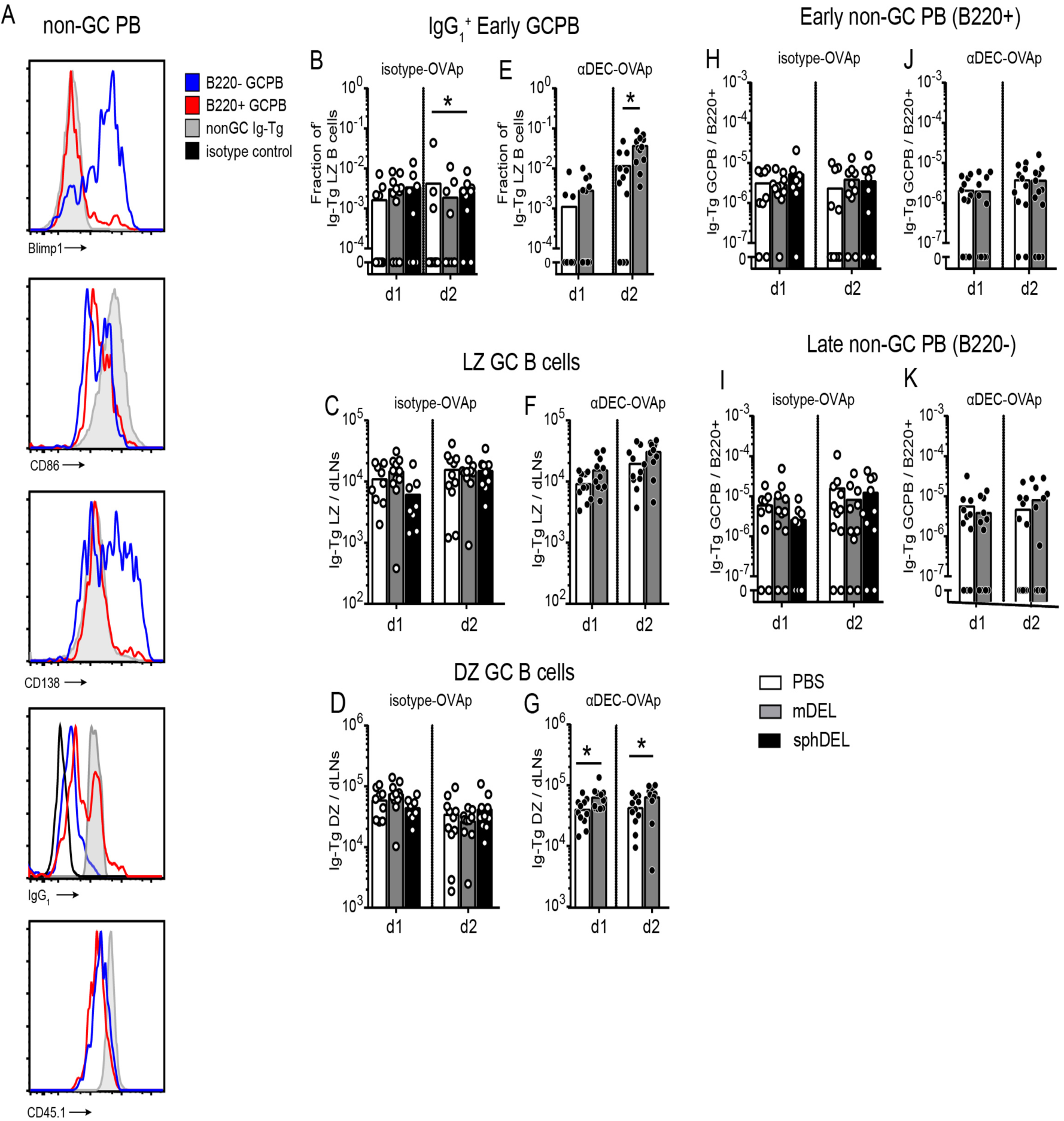
BCR cross-linking enhances early PB differentiation when T cell help is sub-saturating. **A**, representative histographs of Blimp1, CD86, CD138, surface IgG_1_, and CD45.1 in B220^+^(red) and B220^lo^ (blue) GCPBs and non-GC Ig-Tg B cells (grey). For IgG_1_ staining non-GC B cells were gated on IgG_1_^pos^ cells. See **Fig. 3A** for gating strategy. Data representative of 2-3 independent experiments. **B-K**, See **Fig. 3C** for experimental design. Accumulation of IgG_1_^+^ Ig-Tg early (B220^+^) GCPBs as a fraction of IgG_1_^pos^ LZ B cells (in **B, E**). Ig-Tg LZ (**C, F**) and DZ (**D, G**) GC B cells. Early B220^+^ (**H, J**) and late B220 ^lo^ (**I, K**) non-GC Ig-Tg PB in dLNs 1d and 2d post reimmunization. Data from 4 independent experiments. Each symbol represents one mouse. *, p<0.05, Kruskal-Wallis with Dunn’s post test between PBS and mDEL or sphDEL (left panels), or Mann-Whitney test.

